# Super.Complex: A supervised machine learning pipeline for molecular complex detection in protein-interaction networks

**DOI:** 10.1101/2021.06.22.449395

**Authors:** Meghana V. Palukuri, Edward M. Marcotte

**Author notes:** Correspondence to (M.V.P.), (E.M.M.).

## Abstract

Characterization of protein complexes, *i.e.* sets of proteins assembling into a single larger physical entity, is important, as such assemblies play many essential roles in cells such as gene regulation. From networks of protein-protein interactions, potential protein complexes can be identified computationally through the application of community detection methods, which flag groups of entities interacting with each other in certain patterns. Most community detection algorithms tend to be unsupervised and assume that communities are dense network subgraphs, which is not always true, as protein complexes can exhibit diverse network topologies. The few existing supervised machine learning methods are serial and can potentially be improved in terms of accuracy and scalability by using better-suited machine learning models and parallel algorithms. Here, we present Super.Complex, a distributed, supervised AutoML-based pipeline for overlapping community detection in weighted networks. We also propose three new evaluation measures for the outstanding issue of comparing sets of learned and known communities satisfactorily. Super.Complex learns a community fitness function from known communities using an AutoML method and applies this fitness function to detect new communities. A heuristic local search algorithm finds maximally scoring communities, and a parallel implementation can be run on a computer cluster for scaling to large networks. On a yeast protein-interaction network, Super.Complex outperforms 6 other supervised and 4 unsupervised methods. Application of Super.Complex to a human protein-interaction network with ~8k nodes and ~60k edges yields 1,028 protein complexes, with 234 complexes linked to SARS-CoV-2, the COVID-19 virus, with 111 uncharacterized proteins present in 103 learned complexes. Super.Complex is generalizable with the ability to improve results by incorporating domain-specific features. Learned community characteristics can also be transferred from existing applications to detect communities in a new application with no known communities. Code and interactive visualizations of learned human protein complexes are freely available at: https://sites.google.com/view/supercomplex/super-complex-v3-0.

## Introduction

A protein complex is a group of proteins that interact with each other to perform a particular function in a cell, the basic biological unit of all living organisms. Some examples include the elaborate multiprotein complexes of mRNA transcription and elongation helping with gene regulation and key cytoskeletal protein complexes, such as microtubules with their trafficking proteins which help establish major structural elements of cells. Extensive biological experiments have investigated the physical interactions between proteins, and these have been modeled via weighted protein-protein interaction (PPI) networks, where a protein-protein edge weight corresponds to the strength of evidence for the protein-protein interaction. Disruption of protein-protein interactions often leads to disease, therefore identifying a complete list of protein complexes allows us to better understand the association of protein and disease. All experimental protocols for detecting complexes (such as AP/MS, affinity purification with mass spectrometry, and CF/MS, co-fractionation with mass spectrometry) have a tendency to miss interactions (false negatives) and may also predict extra interactions (false positives). Proteins may also participate in more than one complex, potentially blurring the boundaries of otherwise unrelated protein communities. Computational analysis of protein-protein interaction networks can therefore be very useful in identifying accurate protein complexes and will help augment and direct experimental methods.

The weighted PPI network or graph G can be represented as pairs of nodes and edges (*V, E)*, where the set of nodes or vertices *V* represents the proteins, and the set of weighted edges *E* represents the strengths of evidence for interactions between proteins. Any group of nodes and edges that can be characterized as a protein complex can be referred to as a community; community detection methods can be used in turn to identify protein complexes.

A standard guideline for defining communities [1] is that a community should have more interactions or connectivity among the community than with the rest of the network. This can be modeled for example by the community fitness function in Equation 1, mapping a subgraph, *C*, *i.e.* a group of nodes and edges from the full graph, to a scalar value representing a score, where a higher score indicates more community resemblance.

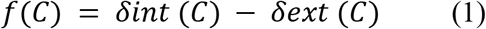

The intra-cluster density *δint* (*C*) and inter-cluster density *δext* (*C*) are given by

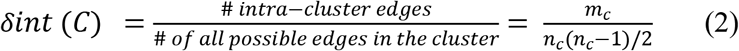

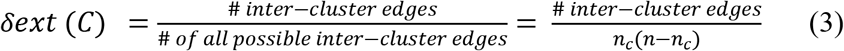

Here, *n*_*c*_ and *m*_*c*_ are the numbers of nodes and edges in subgraph *C,* respectively, and *n* is the number of nodes in graph *G.*

However, there exist many communities that do not follow this criterion but can be identified by different properties they exhibit. One such example is a star-like topology, where one central node interacts with several nodes in yeast protein-interaction networks [2], as, for example, in the case of a molecular chaperone that acts on a number of separate protein clients. In the case of human protein complexes, we also observe different topologies such as clique, linear, and hybrid between linear and clique, as shown **Fig 1**. These human protein complexes represent proteins known to belong to experimentally characterized gold-standard protein complexes from CORUM 3.0 (the comprehensive resource of mammalian protein complexes) [3] with edge weights taken from hu.MAP [4], a human protein interaction network with interactions derived from over 9,000 published mass spectrometry experiments.

**Fig 1.**
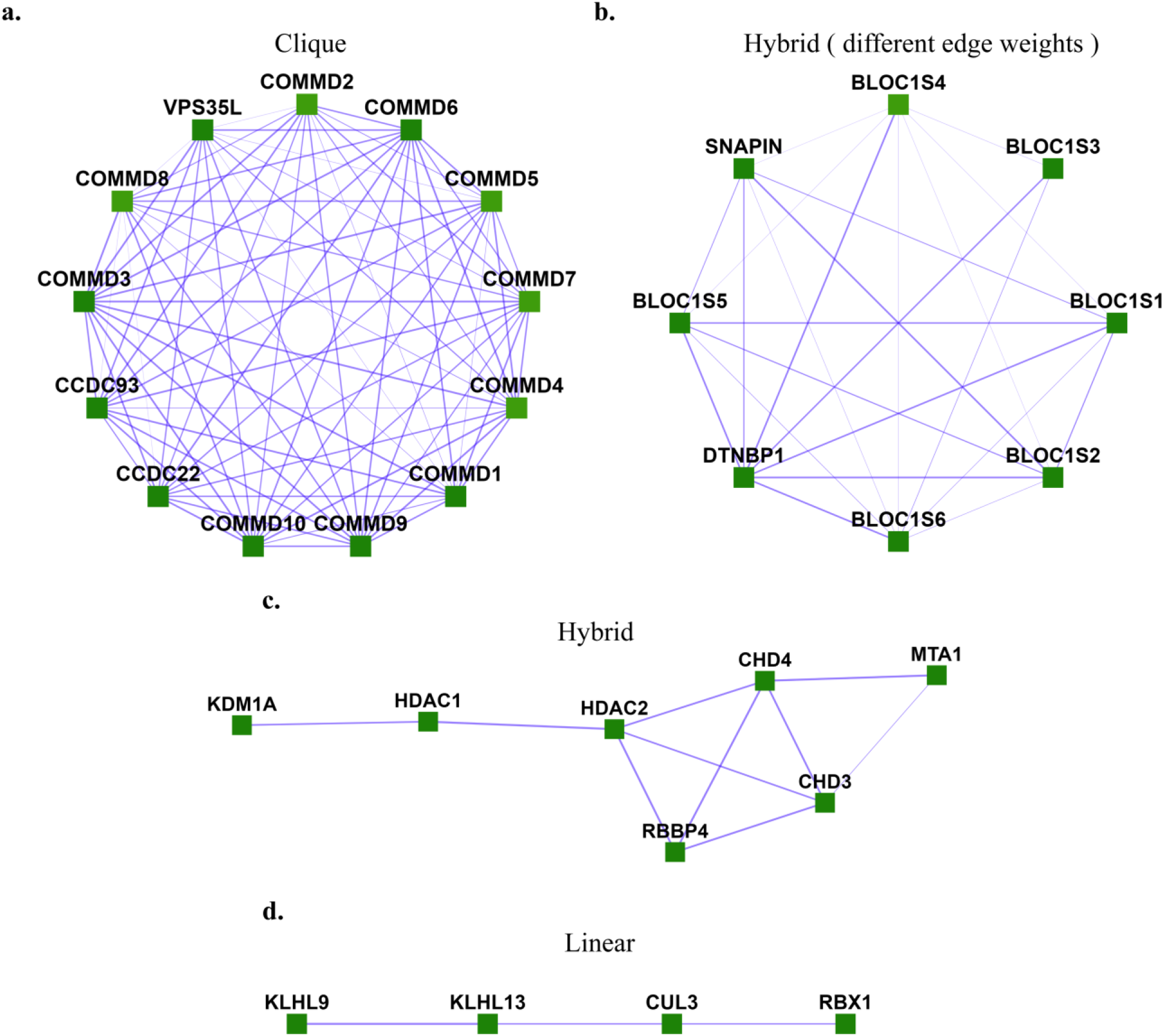
Different topologies are exhibited by human protein complexes. **a.** Clique ([Commander/CCC complex](https://meghanapalukuri.github.io/Complexes/CORUM/Complex112.html)), **b.** Hybrid with different edge-weights ([BLOC-1](https://meghanapalukuri.github.io/Complexes/CORUM/Complex22.html) (biogenesis of lysosome-related organelles complex 1)), c. Hybrid ([NRD complex](https://meghanapalukuri.github.io/Complexes/CORUM/Complex119.html) (Nucleosome remodeling and deacetylation complex), **d.** Linear ([Ubiquitin E3 ligase](https://meghanapalukuri.github.io/Complexes/CORUM/Complex96.html) (CUL3, KLHL9, KLHL13, RBX1)). These are experimentally characterized complexes from CORUM [3] with protein interaction evidence obtained from hu.MAP [4].

Existing community detection methods have primarily tried to optimize for high scores of community fitness functions, such as that of equation 1 [5]. These include unsupervised methods, such as implemented by MCL- Markov Clustering [6], MCODE - Molecular COmplex DEtection [7], CFinder [8], SCAN- Structural Clustering Algorithm for Networks [9], CMC - Clustering based on Maximal Cliques [10], COACH - COre- AttaCHment based method [11], GCE - Greedy Clique Expansion [5], and ClusterONE - clustering with overlapping neighborhood expansion [12], as well as semi-supervised machine learning algorithms such as COCDM - Constrained Overlapping Complex Detection Model [13].

When there are sufficient data available on known communities, rather than applying a generic community fitness function to the problem, it can be more accurate to learn a community fitness function directly from known communities. Then, new communities detected with the learned community fitness function can be expected to better resemble known communities in the field. Supervised machine learning methods are well suited for this purpose, and a few methods have been used to learn a community fitness function from constructed community embeddings, *i.e.*, community representations in vector space, obtained by extracting topological and domain-specific features from communities. The community fitness function learned can then be used to select candidate communities from the network and evaluate them. Since finding maximally scoring communities in a network is an NP-hard (non-deterministic polynomial-time hard) problem [2], heuristic algorithms have been used to find candidate communities. A common strategy is to select a seed (such as a node or a clique) and grow it into a candidate community by iteratively selecting neighbors to add to the current subgraph using heuristics such as iterative simulated annealing until a defined stopping criterion is met for the growth process. This process is repeated with different seeds to generate a set of candidate communities.

Existing supervised methods use different machine learning methods to learn the community fitness function after extracting different features and use different heuristic algorithms to select candidate communities. The first supervised method [2] used a support vector machine (SCI-SVM) and a Bayesian network (SCI-BN) with 33 features with a greedy heuristic, followed by iterative simulated annealing. Stopping criteria for the growth of a seed include limiting the rounds of growth, checking for score improvement over multiple iterations, and checking for overlap with learned candidate communities so far. A second approach [14] recursively trained a two-layer feed-forward neural network model, NN for the classifier using 43 features. This greedy heuristic sequentially grows seeds of the highest degree with similar stopping criteria as [2]. Supervised learning protein complex detection SLPC [15] uses a regression model (RM) with 10 topological features, solved by gradient descent. A modified cliques algorithm finds and grows maximal cliques using a random but exhaustive neighbor selection followed by a greedy growth heuristic. The algorithm stops when no node addition can yield a higher score, after which they merge some pairs of overlapping complexes with an overlap greater than a threshold. ClusterEPs, short for cluster emerging patterns [10] uses a score function based on noise-tolerant emerging patterns (NEPs) which are minimal discriminatory feature sets using 22 features, along with an average node degree term. Like [14], the heuristic for this method also grows the highest degree seed nodes sequentially. The neighboring node that shares the maximum number of edges with the current subgraph is selected as a candidate for growth in each iteration and a greedy growth heuristic is used, stopping when the score is greater than 0.5. ClusterSS, short for clustering with supervised and structural information [16] uses a neural network with one hidden layer and 17 features, along with a traditional structural score function from [12]. A greedy heuristic grows seed nodes, also considering deletion of any existing subgraph nodes, with an optimization step of considering only the top k nodes by degree. The stopping criterion is when the new score is less than a factor times the old score. Both ClusterEPs and ClusterSS merge pairs of communities with overlap greater than a threshold at the end.

Regarding scalability, the above methods have generally only been implemented on small yeast protein complex datasets, except for ClusterEPs, which trains on yeast data and tests on human PPIs. [17] implement the regression model of [15] on a human PPI network re-weighted by breast-cancer specific PPIs extracted from biomedical literature to detect disease-specific complexes. However, these methods employ serial candidate community sampling, negatively impacting their scalability to large networks such as hu.MAP [4], a human protein-interaction network with ~8k nodes and ~60k edges.

In this work, we present Super.Complex (short for Supervised Complex detection algorithm), an end-to-end highly scalable (to large networks that fit on a disk), distributed, and efficient community detection pipeline that explores multiple supervised learning methods with AutoML (Automated Machine Learning) to learn the most accurate community fitness function from known communities. Super.Complex then samples candidate subgraphs in parallel by seeding nodes or starting with maximal cliques and growing them with an epsilon-greedy heuristic, followed by an additional heuristic such as iterative simulated annealing or pseudo-metropolis using the learned community fitness function. On a yeast PPI network, Super.Complex outperforms all 6 existing supervised methods, as well as 4 unsupervised methods. Three novel evaluation measures are proposed to overcome certain shortcomings of existing metrics. We apply Super.Complex to hu.MAP, a human protein-protein interaction network with ~8k nodes and ~60k edges to yield 1028 protein complexes, including high-scoring previously unknown protein complexes, potentially contributing to new biology, and make all data, code, and interactive visualizations openly and freely available at https://sites.google.com/view/supercomplex/super-complex-v3-0.

## Materials and Methods

### Overview of Super.Complex

The pipeline Super.Complex comprises two main tasks, first, learning a community fitness function with AutoML methods, and second, using the community fitness function to intelligently sample overlapping communities from a network in parallel. As shown in **Fig 2**, each task is subdivided into different steps, described in brief in this section, with all details in the following sections of Materials and Methods. For the first task, we perform a pre-processing step, Data Preparation, where known communities are cleaned and split into non-overlapping training and testing sets, followed by construction of training and testing negative community data. In (i) Topological Feature Extraction, topological characteristics for all communities are computed to construct training and testing feature matrices. AutoML (ii) then compares different ML (Machine Learning) pipelines to select the best one, followed by training and testing the best ML pipeline, thus learning the community fitness function as the binary classifier distinguishing positive communities from negatives. Having learned the community fitness function, Super.Complex then uses it in its heuristic algorithm for the second task of searching for candidate communities in the network in parallel. For (iii) intelligent sampling, the algorithm can start with either single nodes or maximal cliques as seeds. We note that all nodes of the network were used as seeds in our experiments (this is quite fast due to Super.Complex’s parallel implementation), allowing us to work without any estimate of the number of expected communities. These seeds are grown using a 2-stage heuristic, 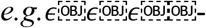greedy + iterative simulated annealing. This is followed by a (iv) post-processing step of merging highly overlapping communities. Finally, in the last step, evaluation, the learned communities are compared with known communities. The steps of the pipeline are fairly independent and can be improved on their own with methods to test the accuracy/performance of each of the steps.

**Fig 2.**
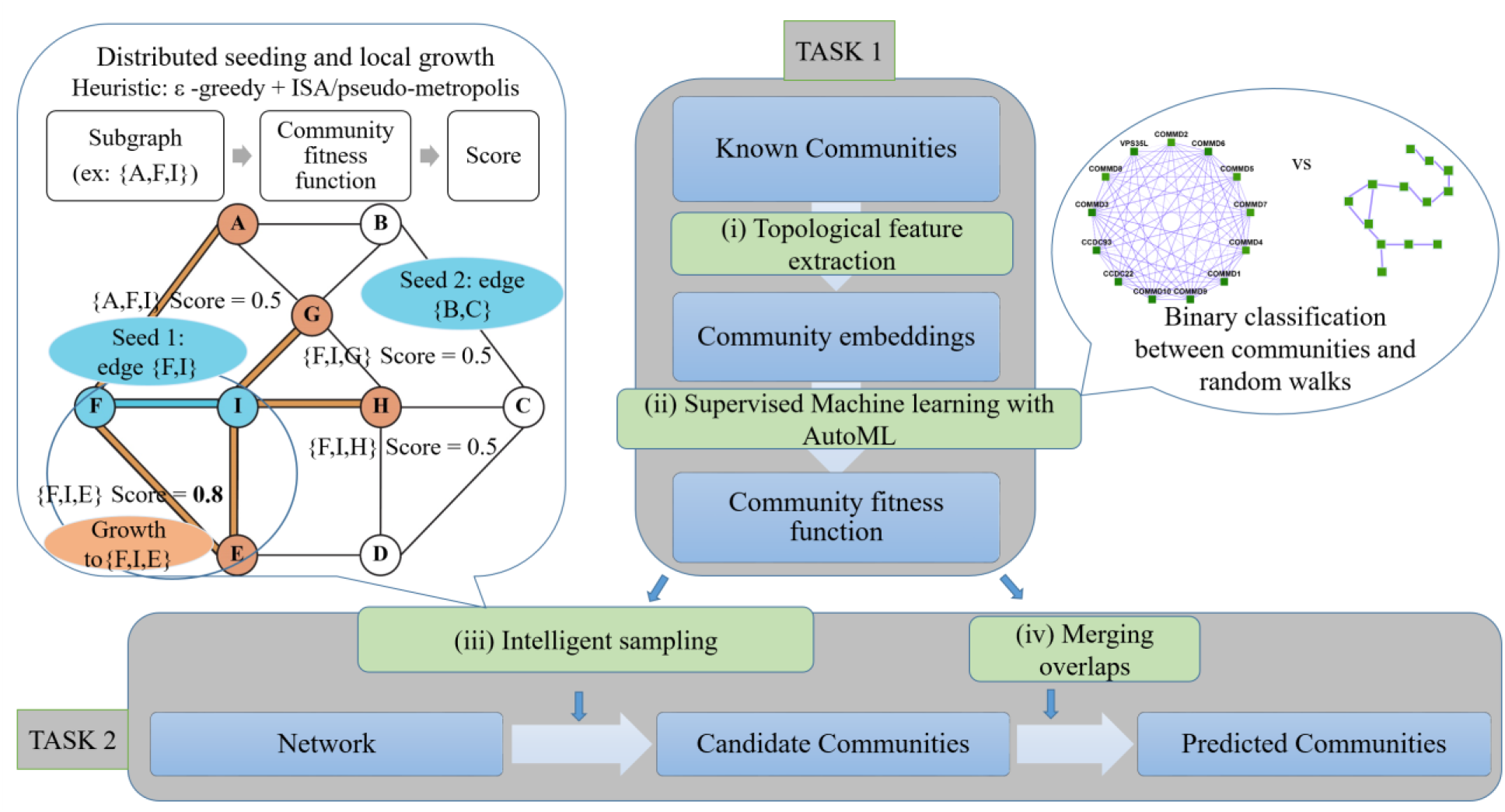
Super.Complex identifies likely protein complexes within a PPI network using a distributed supervised AutoML method. Task 1: Learning a community fitness function: (i) Topological feature extraction: Topological features are extracted from known communities to build community embeddings (feature vectors, which are representations of communities in vector space) (ii) Supervised learning with AutoML: A score function for communities, the community fitness function, is learned from the community embeddings as the decision function for binary classification of a network subgraph as a community or a random walk (illustration on the right). The best score function is selected after training multiple machine learning models with TPOT [18], an AutoML pipeline. Task 2: Searching for candidate communities in the network: (iii) Intelligent sampling: Multiple communities are sampled in parallel from the network. To build each candidate community, a seed edge is selected and grown using a 2-stage heuristic. First, we use an epsilon-greedy heuristic to select a candidate neighbor, and then we use a pseudo-metropolis (constant probability) or iterative simulated annealing heuristic to accept or reject the candidate neighbor for growing the current community. An iteration of neighbor selection using a greedy heuristic is shown (illustration on the left), starting from a seed edge {F, I}. The edge is grown to the subgraph {F, I, E} as adding node E yields a higher community fitness function than adding any other neighbor of F and I. The seed edge {B, C} is grown in parallel (not shown) (iv) Merging overlaps: The candidate communities are merged such that the maximum overlap between any 2 communities is not greater than a specified threshold.

### Data Preparation

First, the weighted network under consideration is cleaned by removing self-loops, as we do not consider interactions with oneself as a feature of communities. For scalability, the graph is stored on disk as a set of files, each corresponding to a node and containing a list of the node’s neighbors via weighted edges.

#### Positive communities

Super.Complex takes sets of nodes comprising known communities and obtains their edge information from the induced subgraph of these nodes on the weighted network. Nodes in communities that are absent from the network are removed. Communities with fewer than 3 nodes, communities that are internally disconnected, and duplicate communities are also removed. Constructing the final set of positive communities involves 2 main steps: (i) merging similar communities, and (ii) splitting them into non-overlapping train and test sets. Note that if independent train and test sets of communities are known in advance, these steps can be skipped.

In the first step, using a merging algorithm we devised, we merge highly similar communities to yield a final list of communities, where no pair of communities have a Jaccard score (**S1 File** equation 3) greater than or equal to *j.* We recommend users to set this value based on domain knowledge of observed redundancy in the set of known communities.

Multiple solutions exist that achieve this goal, however, we want a solution with a large number of communities, *i.e.*, with only a small number of merges performed on the original set of known communities. This is especially important in applications with limited data, such as the human and yeast protein complex experiments in this work. Our algorithm was designed with this objective in mind, and works as follows. The iterative algorithm makes multiple passes through the list of communities performing the merging operation until the specified criterion is achieved. In a single pass of the list of communities, each community is considered in order and merged with the community with which it has the highest overlap (if greater than or equal to *j*) and the list is updated immediately by removing the original 2 communities and adding the merged community to the end of the list, so that the updated list is available for the next community in consideration. This merging algorithm achieves a lesser number of merges than a trivial merging solution which would merge random pairs of communities that do not satisfy the required criteria until convergence. In practice, the proposed algorithm quickly converges to a solution (*i.e.* the final set has no communities that overlap more than the specified value *j*).

In the second step, the communities are split into non-overlapping training and testing datasets, to emphasize their independence. We obtain sets with equal size distributions and a 70-30 train-test split, as recommended for machine learning algorithms with a small amount of data. Previous algorithms such as [4] and Super.Complex v2.0 [31] discard test communities with sizes greater than a threshold, thus losing out on information from some known communities and which also, in practice, do not yield train-test splits that are close to the recommended 70-30 split. Therefore, we propose the following algorithm. Here, we first make the recommended 70-30 random split into train and test communities. Then we perform iterations of transfers between the two sets until they become independent. In each iteration, we perform two directions of transfers, from train to test and vice-versa, and if the 70-30 split is disturbed, we remove the communities at the end of the list which have extra communities and add them to the other list. In each direction of transfer, for instance, from train to test, we go through the training communities in one pass and if a training community has an overlap (at least one edge) with any of the test communities, it is immediately transferred to the test set, making the updated test set available for comparison with subsequent training set communities. In practice, for many random splits, the algorithm converges fast enough to a solution that is non-overlapping. If for an initial random split, convergence is not achieved after a few iterations, we recommend restarting the algorithm with a different random split.

#### Negative communities

Negative communities, or non-communities are represented by random walks sampled from the network by growing random seeds, adding a random neighbor at each step. The number of steps ranges from the minimum size to the maximum size of positive communities, with a total number of random walks equal to the number of positive communities multiplied by a scale factor > 1. The random walks are split almost equally across all the sizes, by splitting equally across the different number of steps to be taken for a random walk, to yield an almost uniform size distribution for negative communities. We say almost uniform size distribution, as random walks with the same number of steps need not yield the same sizes, given that the random walk as defined here can revisit edges it has already visited. To achieve random walks of the same size, the algorithm attempts an extra number of random walks and an extra number of steps to achieve the desired random walk size.

The size distribution of positive communities is taken into consideration while training the machine learning model when using a uniform distribution for negatives. We also explore using almost the same size distribution as the positive communities to construct the negative communities. For this, for each size of the positive communities, we construct the negatives by sampling a number of random walks equal to the scale factor times the number of positive communities of this size. However, in this case, we find that there are quite a few missing sizes due to limited positives which may affect the scoring of subgraphs of the missing sizes. Using a uniform distribution would provide more information to learn a more accurate community fitness function that can recognize negatives at sizes missing for positives. In the following feature extraction step, random walks resembling communities are removed. The final number of negative communities is close to the number of positive communities, as we have sampled a slightly higher number of random walks via the scale factor.

### Topological Feature Extraction

As communities exhibit different topological structures on the graph, these can be learned by considering useful topological features of communities. Based on graph theory, we extract 18 topological features, detailed in **S1 File** Methods (Topological features) for each of the positive/negative communities to construct the final train and test data feature matrices, *i.e.* the positive and negative community embeddings.

### Learning the community fitness function with AutoML

A community fitness function is learned as the decision function of a binary machine learning classifier trained to distinguish the community and non-community embeddings constructed in the previous feature extraction step. For this, we use an AutoML algorithm, TPOT [18], a genetic algorithm that yields the best model and parameters. It evaluates several preprocessors along with ML models and yields cross-validation scores on the training dataset for each pipeline, which itself is usually a combination of several preprocessors followed by the machine learning model. We configure the algorithm to run in a distributed setting, exploring several combinations of several preprocessors and ML models.

We specify 6 pre-processors that scale the feature matrix. These are - (i) Binarizer, which sets a feature to 0 or 1 based on a threshold, (ii) MaxAbsScaler, which divides the feature by the maximum absolute value of the feature, (iii) MinMaxScaler, which subtracts the minimum of the feature from the feature vector and divides by the range of the feature, (iv) Normalizer, which divides the feature vector by its norm to get a unit norm, (v) RobustScaler, which makes a feature robust to outliers by scaling using the interquartile range and (vi) StandardScaler, which standardizes to the Z-score by subtracting the mean and dividing by the standard deviation of the feature.

We include four feature selecting pre-processors, which are additionally important as we incorporate 6 additional preprocessors that construct combined features. The additional preprocessors include - (i) Decomposition: PCA (Principal Component Analysis), FastICA (Independent Component Analysis), (ii) Feature Agglomeration, (iii) Kernel Approximation methods: Nystroem, Radial Basis Function RBFSampler, (iv) Adding Polynomial Features, (v) Zero counts: Adds the count of zeros and non-zeros per sample as features and (vi) OneHotEncoder for numeric categorical variables. The feature selecting preprocessors include - (i) SelectPercentile, which selects the highest-scoring percentage of features based on 3 univariate statistical tests, FPR - False Positive Rate, FDR - False Discovery Rate and FWE - Family-wise error rate; (ii) VarianceThreshold which removes low variance features, (iii) RFE (recursive feature elimination) using ExtraTrees and (iv) SelectFromModel using ExtraTrees based on importance weights. The ML models included are - (i) Naive Bayes methods using Gaussian, Bernoulli, and Multinomial distributions (ii) Decision Trees, (iii) Ensemble methods of ExtraTrees, Random Forest, Gradient Boosting and XGB (XGBoost), (iv) K-nearest neighbors, (v) Linear SVMs and (vi) Linear models for Logistic Regression.

The population size and number of generations are provided as parameters for the genetic algorithm of the AutoML pipeline. In practice for our application, giving a value of 50 for each yielded good results. There is an option for a warm start, where you can run additional generations and with additional population sizes starting from the latest results, if the results are unsatisfactory. Additionally, several other machine learning models and preprocessors can also be incorporated into this pipeline, including neural networks. Note that in our experiments, we also obtained pipelines that stack different ML models. We run the pipeline in a distributed manner, setting the number of jobs as the number of processes that run in parallel on a single computer. All the processes on the computer can be used for maximum utilization, however, the documentation notes that memory issues may arise for large datasets. In practice, we set the number of jobs as 20 on a Skylake compute node (Intel Xeon Platinum 8160 with 48 cores @2GHz clock rate).

#### Evaluation

By default, 5-fold cross-validation is performed, although this can be modified by a parameter. The pipelines with high cross-validation average precision scores (area under the PR curve) are evaluated on the test dataset to find the best pipeline for our data, to use this for the community fitness function. A one hidden-layer perceptron is also available for training, and comparison with the AutoML output to select the best model. We evaluate the performance of the ML binary classifier using accuracies, precision-recall-f1 score measures, average precision score, and PR curves for the test sets while also evaluating these measures for the training set to compare with the test measures and check the bias and variance of the algorithm to make sure it is not underfitting or overfitting the data. We also plot the size-wise accuracies of the model to understand how a model performs w.r.t to the size of the subgraph it is evaluating.

### Candidate community search

Finding a set of maximally scoring candidate communities in a network is an NP-hard problem, as proved by [2] by reducing it to the problem of finding maximal cliques. Since this is an NP-hard problem, algorithms based on heuristics are required to solve it. We explore seeding and growth strategies.

#### Design and distributed architecture

First, we need to select seeds. Options for seeds include specifying all the nodes of the graph (recommended for best accuracy), all the nodes of the graph present in known communities, a specified number of nodes that will be selected randomly from the graph, or maximal cliques. In the distributed setting using multiple compute nodes, the specified seeds are partitioned equally across compute nodes, and each compute node deals only with the task of growing the seeds assigned to it. In practice, the partitioning is done by a main compute node which partitions the list of seeds and stores the partitioned lists as separate files on the file server. Then it launches one task per compute node (including itself) using the launcher module [32], where a task instructs a compute node to read its respective file containing the seed nodes and run the sampling algorithm starting with each of the seed nodes. On each compute node, we take advantage of all the cores by employing multiprocessing with the *joblib* python library. Each process intelligently grows a single seed node into a candidate community and writes it to the compute node’s temporary storage. For this, we need the graph and parameters of the community fitness function, which we store on temporary disk space of each compute node to optimize RAM as it is impractical to store large networks and machine learning models in memory. Each process reads the model into its memory and uses it to evaluate the neighbors, to pick the neighbor to add to the current subgraph in the growth process from the seed node. The neighbors of the subgraph under consideration at each step of the growth are read from disk on-demand and stored in memory only until they have been evaluated by the fitness function. In this way, we ensure that the processes have a low memory footprint, which can otherwise quickly become a bottleneck for large graphs. We also minimize disk storage by storing each resulting candidate community compactly using only its nodes, as its edges can be inferred if/when necessary by inducing the nodes on the graph. After all the child processes of growing seeds complete on a compute node, the compute node reads the set of learned community files it had stored on its disk and compiles them into a list of candidate communities before writing the list to the file server. The same code also runs in a distributed setting with only one multi-core compute node. There also exists a serial option to run the code without invoking parallel constructs, useful for running on a single core.

#### Intelligent sampling - Heuristics

Only for the first step of growth, we add the neighbor connected with the highest edge-weight. We provide 2 options for growing the subgraph at each step- an exhaustive neighbor search that is suitable for graphs that are not very large, and an option that optimizes performance by evaluating only a subset of neighbors. In the latter, using a large user-defined threshold *t*_1_, if the number of neighbors of the current subgraph is greater than the threshold, a random sample of the neighbors equal to the provided threshold is chosen for evaluation. Now, of the neighbors, first, an *ϵ*-greedy heuristic is used to select the neighbor to add to the subgraph. In an *ϵ*- greedy heuristic, with *ϵ* probability, a random neighbor is added instead of the maximum scoring neighbor.

In the non-exhaustive search case, in the event of 1-*ϵ* probability, if the number of neighbors is greater than a 2nd user-defined threshold *t*_2_, a 2nd optimization of cutting down the number of neighbors is applied before evaluating each of the neighbors for choosing the greedy neighbor, as follows. Here, the *t*_2_highest neighbors are chosen for evaluation, where the order is decided by sorting the neighbors in descending order based on their maximum edge weight (*i.e.* the highest edge weight among all the edges connecting a neighbor to the subgraph). Note how the first threshold *t*_1_ensures that the sorting complexity *0*(*t*_1_*log*(*t*_1_))does not blow up.

Note that for efficient constant-time *0*(1) lookup of the maximum edge weight of a neighbor, we store the neighbors of the subgraph as a hash map, where looking up a neighbor yields its maximum edge weight. This hash map also stores, for each neighbor, a list of edges connecting it to the subgraph and was constructed efficiently when the neighbors of each of the subgraph nodes were read from the corresponding file. After selecting the neighbor to add to the subgraph in the current iteration, this hash map is also used to efficiently add the neighbor to the subgraph by providing constant-time lookup to the edges that need to be added.

Instead of the base *ϵ*-greedy heuristic, we also have a simple base heuristic option, termed greedy edge weight, where we add the neighbor with the highest maximum weight edge at each step of the iteration. Note that since the ML model is not used at each stage of the growth, this is fast enough and does not require the optimization steps used in the *ϵ*- greedy approach where subsets of neighbors were selected for evaluation by the community fitness function.

For both base heuristics, in any iteration, if no neighbors for the subgraph exist, the growth process terminates. If the community score of the subgraph in any iteration is less than 0.5, the node last added is removed and the growth process terminates. We provide additional heuristics that can be applied on top of the base *ϵ*-greedy heuristic. Based on the scores of the current and previous iterations of the subgraph, we accept or reject the latest node addition using the user-defined heuristic - iterative simulated annealing (ISA), or a variant of ISA, termed pseudo-metropolis in which the acceptance probability (equation 9) is a constant, *i.e. P*(*S*_*new*_, *S*_*old*_) = *k*. In ISA, at each stage of growth of the current subgraph, its maximum scoring neighbor is added, except in the case when the new community score of the subgraph *S*_*new*_is lesser than *S*_*old*_, the value before adding the new node (*i.e. S*_*new*_ < *S*_*old*_). In this case, the new node addition is accepted with a probability of,

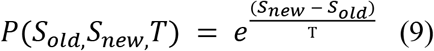

here, starting with hyperparameters *T*_*0*_ and *α*, we update the temperature as *T* ← *αT* after every iteration.

When ISA or pseudo-metropolis heuristics are applied, we also evaluate an additional heuristic where the algorithm terminates if it has been 10 (or can be user-defined) number of iterations since the score of the subgraph has increased.

In the implementation, we provide four options to the user - greedy edge weight,*ϵ*-greedy, *ϵ*-greedy + ISA and *ϵ*-greedy + pseudo-metropolis. In all options, the algorithm terminates after a number of steps equal to a user-specified threshold. The default threshold provided is the maximum size of the known communities, and we also provide a smart option for when a few communities have a large number of nodes, where it is set to choose the maximum size after ignoring outliers. This number can also be improved by visual inspection of a boxplot of community sizes that is generated. Future work can also explore greedy edge weight + ISA and greedy edge weight + pseudo-metropolis heuristic algorithms and observe their performance. Note how there are 2 possibilities for exploration in the 3 algorithms other than the greedy edge weight heuristic algorithm. In the 1st stage, we pick a neighbor at random with low probability. In the 2nd stage, we accept the neighbor we picked in the 1st stage with low probability, if it yields a lower score than the original subgraph.

#### Post-processing (merging overlaps) and cross-validation

Communities with only 2 nodes are removed. Note that communities with 2 nodes are rarely found, and while dimers are biologically valid, since they do not have topological variation, we do not consider them in this work focused on higher order assemblies with different topologies as a key feature. We then merge communities that have a Jaccard similarity greater than a specified overlap threshold employing the merging algorithm discussed in the data cleaning section. The only difference is while merging, for two overlapping communities, the final community retained out of the 2 communities or the merged variant is the one that obtains the highest score with the community fitness function. In another variant of the merging algorithm, instead of the Jaccard similarity threshold (**S1 File** equation 3), we use Qi’s overlap measure (**S1 File** equation 10).

The parameters *ϵ*in the *ϵ*-greedy heuristic, k in pseudo-metropolis, *T*_*0*_and *α*in iterative simulated annealing, and the overlapping threshold in the post-processing step are varied in parameter sweeps to select the best ones that work using the Qi et al F1 score (**S1 File** equation 8).

After parameter sweeps, the results of different heuristics are examined and the one that yields the best F1 score is chosen. Additional details regarding evaluation are outlined in S1 File Methods (Evaluation with existing measures).

## Results and Discussion

### Contributions of Super.Complex - a scalable, distributed supervised AutoML-based community detection method

Super.Complex implements an original distributed architecture and an efficient pipeline, scaling to large networks such as hu.MAP with ~8k nodes and ~60k edges. With an AutoML method, which also includes automated feature selection, and four 2-stage heuristic options for candidate community search, the pipeline finds accurate community fitness functions and high quality communities. Unlike some existing methods that remove nodes in the process of growth (e.g. such as Louvain [19] and ClusterSS), we note in **S1 File** Results (Algorithm guarantees) that our method guarantees properties such as internal connectivity of communities. Further, the merging algorithm we employ guarantees that no two communities overlap more than a specified threshold. In the case of non-overlapping communities (obtained by specifying a merging threshold of 0 overlap), there is an additional guarantee that no two communities can be merged to yield a higher scoring community. To our knowledge, epsilon-greedy heuristics in conjunction with other heuristics such as iterative simulated annealing have not been applied in the past for community detection. This allows the pipeline to leverage advantages of both heuristics by adding an additional layer of stochasticity allowing better exploration in the candidate community search stage. Super.Complex has a cross-validation pipeline to select the heuristic and parameters that work best for the application at hand. Minimal hyper-parameter selection is required in our algorithm with default parameters provided when smart hyperparameters cannot be inferred.

Since the number of known communities can be limited, we emphasize the preservation of known communities when splitting them into train-test sets while also ensuring (i) independence - *i.e*., no edge overlap between a train and test community on the network, (ii) similar size distributions for both sets, and (iii) 70-30 ratios in train-test sets. Similarly, a minimal number of merges is attempted in the merging algorithms devised to maintain a high number of learned protein complexes. Further, unlike existing supervised methods, which evaluate the performance of their algorithms on a reduced network with only nodes present in known communities, we evaluate our algorithm on the full network for more accurate evaluation. Finally, we note that Super.Complex uses only topological features of networks, and can be applied to community detection on networks from various fields, with the possibility of including domain-specific features to learn more accurate domain-specific community fitness functions. Our methods are also applicable in domains with limited or no knowledge by transferring community fitness functions from other domains, such as the defaults we provide for human protein complex detection.

### Three novel evaluation measures to compare learned communities with known communities

Comparing sets of learned and known communities accurately is an outstanding issue. Poor evaluation measures do not satisfactorily identify the quality of learned communities and make it difficult to evaluate a community detection algorithm. Sets of learned communities achieving high scores with existing evaluation measures have been observed to have a lot of redundancies, e.g. multiple learned communities are very similar with high overlaps [20]. Known big communities were also observed to be split into several learned communities while still achieving good scores on evaluation measures. While it is undesirable to have many false negatives, having many false positives is more hurtful, as wet-lab experiments for biological validation tend to be quite expensive and time-consuming to perform. Therefore, we concentrate on including precision-like measures that compute false positives. Further, evaluation measures that are not sensitive to changes in the sets of learned communities limit our abilities to iterate successfully over algorithm modifications to improve algorithms. We examine the specific shortcomings of different evaluation measures and propose new measures to help overcome the issues discussed and construct robust yet sensitive measures.

#### F-similarity-based Maximal Matching F-score (FMMF)

An issue with many measures such as Qi et al F1 score (**S1 File** equation 8) and SPA (**S1 File** equation 9) is that they don’t penalize redundancy, *i.e.* if we learn multiple same or very similar communities which are each individually high scoring, we will get a high value of precision-like measures. This is because in many cases, many to one matches are being made between learned communities and known communities. To deal with such issues, it is best to make one-to-one matches. The MMR (Maximal Matching Ratio) is one such good measure, however, it only calculates a recall-like measure by dividing the sum of the weights of edges (in a maximal sum of one-to-one edge weights) by the total number of known communities. Taken alone this cannot account for precision, for instance, if we learn a series of random subgraphs, these have low weights and will be ignored, while a high MMR score can be obtained from only a small number of high quality learned communities. Therefore, we define the precision equivalent for MMR, *P*_*FFM*_ in **Fig 3c**.

**Fig 3.**
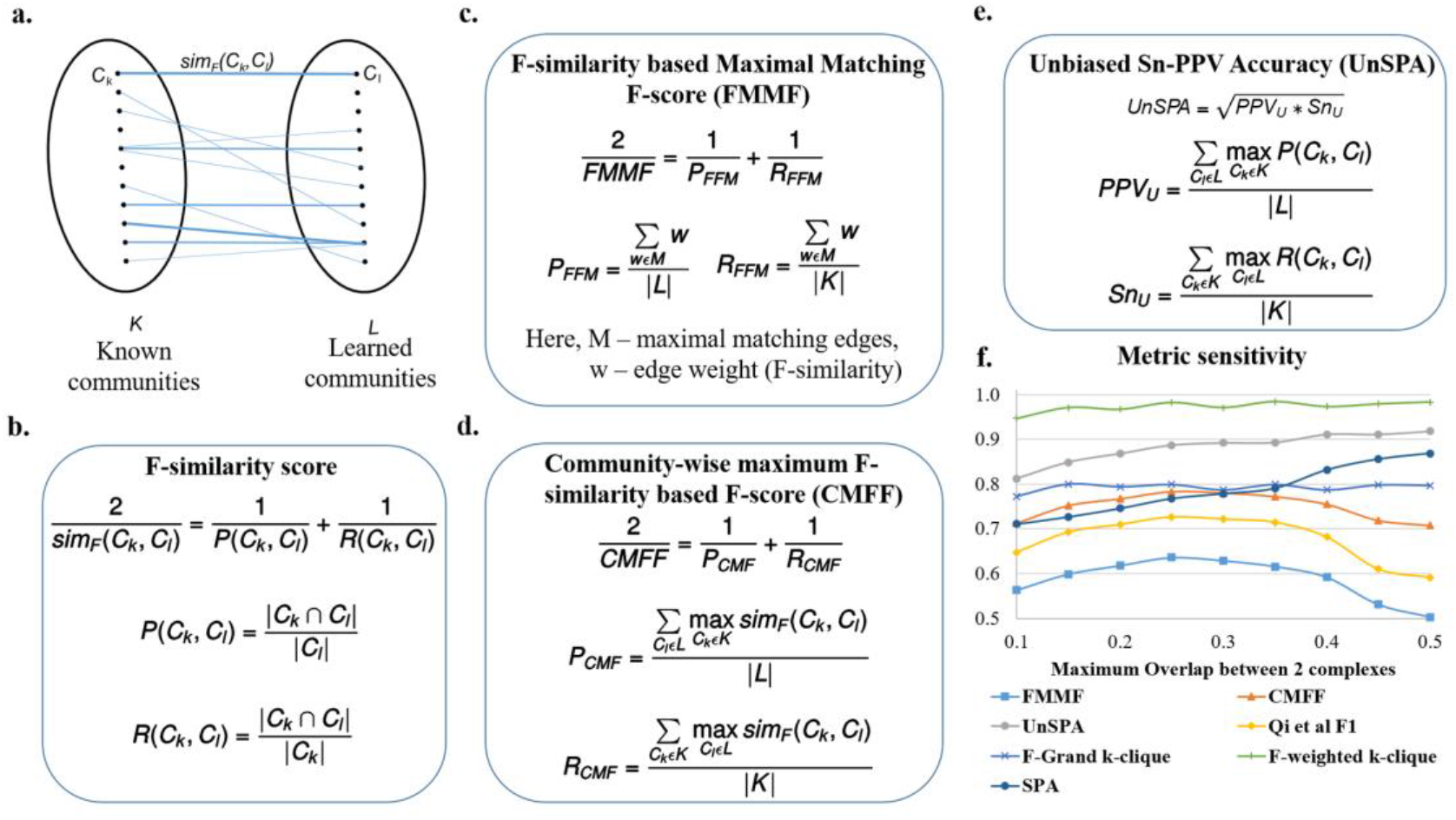
Proposed evaluation measures - FMMF, CMFF, and UnSPA are sensitive metrics. **a.** Bipartite graph, where each edge weight corresponds to the F-similarity (*sim*_*F*_(*C*_*k*_, *C*_*l*_)) between *C*_*k*_, a known community from *K*, the set of known communities and *C*_*l*_, a learned community from *L*, the set of learned communities. **b**. The F-similarity score combines precision (*P*(*C*_*k*_, *C*_*l*_)) and recall (*R*(*C*_*k*_, *C*_*l*_)) measures, computed as fractions of the number of common nodes w.r.t the number of nodes in a community. |*C*|is the number of nodes in community *C* and | *C*_1_ ∩ *C*_2_ | is the number of nodes common to both communities. **c.** F-similarity-based Maximal Matching F-score (FMMF) combines precision (*P*_*FFM*_) and recall (*R*_*FFM*_) measures computed for a maximal matching, *M* of the bipartite graph in **Fig 3a d.** Community-wise Maximum F-similarity based F-score (CMFF) combines precision (*P*_*CMF*_) and recall (*R*_*CMF*_) measures, averaging over the maximum F-similarity score for a community in a particular set (e.g. known communities) w.r.t to a community of the other set (e.g. learned communities) **e.** UnSPA is an unbiased version of Sn-PPV accuracy (SPA), computed as the geometric mean of unbiased PPV (*PPV*_*u*_) and unbiased Sensitivity (*Sn*_*u*_), computed similar to precision and recall measures in CMFF, only, instead of the F-similarity score, precision and recall similarity scores are used respectively **f.** Sensitivity of different evaluation measures w.r.t. (maximum pairwise Jaccard coefficient) overlap between communities shows that FMMF, CMFF, UnSPA, and existing measures Qi et al F1 score (**S1 File** equation 8), and SPA (**S1 File** equation 9) are sensitive metrics, with FMMF, CMFF, and Qi et al F1 score following the desired trend. Here, each data point on the plot corresponds to a measure evaluating an individual run of Super.Complex’s merging algorithm with a maximum Jaccard overlap threshold set to the x-axis value.

In **Fig 3c**, *M* is a set of weights of a set of maximal one-to-one matches, found using Karp’s algorithm [21]. The weight w that we use is the F-similarity score (**Fig 3b**), also described in the next section, Community-wise Maximum F-similarity based F-score (CMFF), unlike the neighborhood affinity used in the original MMR. Correspondingly we can define an F-score, FMMF, as the harmonic mean of the precision *P*_*FFM*_and recall *R*_*FFM*_, also shown in **Fig 3c**.

By doing a one-to-one match, we are also indirectly penalizing cases where the benchmark community is split into multiple smaller communities in the learned set of communities, since the measure considers the weight of only one of the smaller learned communities that comprise the known community, ignoring the rest. Thus only the small weight of the matched community is considered, penalizing this case, unlike one-to-many measures that aggregate the contributions from each of the smaller communities to finally achieve a high score.

#### Community-wise Maximum F-similarity-based F-score (CMFF)

[5] compute F1 scores at the individual known community-learned community match level and look at the histograms of these scores for all known communities. While their work does not state the exact formulation of their F1 score, we are inspired by them to define an F1 score at the match level, *i.e.* an F-similarity score, by comparing the nodes of a learned and a known community. Our F-similarity score is a combination of the recall (of the nodes of the known community) and the precision (of the nodes of the learned community), as shown in **Fig 3b**.

Our F-similarity score can be compared with a threshold to determine a match, and then the overall precision, recall, and F1 scores for the set of predictions can be computed as in **S1 File** equations 6-8. Alternatively, our F-similarity score can be used to determine the best matches for communities and overall measures can be defined that can be investigated to reveal the contributions at the individual match level as well. For interpretability at the match level, similar to the unbiased sensitivity and PPV metrics (as discussed in the next section, Unbiased Sn-PPV Accuracy (UnSPA)), we can define precision and recall measures that evaluate, for each community, the closest matching community in the other set using a similarity metric. Using the F-similarity score as the similarity metric here, we define precision and recall-like measures, and combine them into the F1-like measure, CMFF - Community-wise Maximum F-similarity based F-score, as shown in **Fig 3d**. We detail a general framework to construct similar measures in the next paragraph, drawing inspiration from modifications to the Qi et al F1 score. This framework also gives another method of constructing the CMFF.

In the Qi et al measures from [2], (**S1 File** equations 6-8), a binary indication of a possible match is used, *i.e.* as long as there exists a possible match, it is used as a 1 or 0 count towards the aggregate precision or recall measures. Having a measure that provides matches between learned and known communities allows easy identification of previously unknown communities. One to many matches such as Qi et al precision-recall (PR) measures that do not use an explicit matching between learned and known communities can be modified to obtain a matching. In the modified measure, for each community, we choose the most similar community in the other set in order to give the matching. While measures that use a threshold such as Qi et al F1 score (**S1 File** equation 8) have the advantage of being robust, until a match crosses a threshold, the measure will not change, making it insensitive to small variations in predictions. Measures with low sensitivity make it difficult to compare algorithms and select parameters. Weighted measures are more sensitive, giving different values based on the quality of matches, and are more precise when compared to summing binary values of match existence. Accordingly, a more sensitive and precise version of the Qi et al F1 score can be obtained by summing up weights indicating the similarity scores. For instance, instead of the Qi overlap measure (**S1 File** equation 8), the neighborhood affinity similarity measure (**S1 File** equation 4) can be used to construct a more precise and sensitive measure.

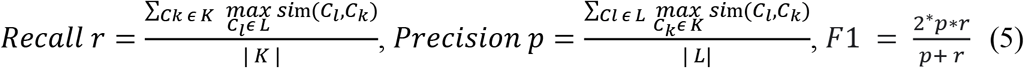

Here, *sim*(*C*_1_, *C*_2_)is a similarity measure between communities *C*_1_and *C*_2_, with |*C*_1_|is the number of nodes in *C*_1_. *C*_*k*_ is a known community from *K*, the set of known communities and *C*_*l*_is a learned community from *L*, the set of learned communities.

Different similarity measures (**S1 File** equations 3–5), such as the Jaccard coefficient can be used to construct different F1 measures. We recommend the F-similarity measure in **Fig 3b**, as it can be broken down into a precision-based and recall-based measure at the level of comparing a known and learned community, and use it to construct the CMFF score.

#### Unbiased Sn-PPV Accuracy (UnSPA)

Consider the precision-like positive-predictive value (PPV), recall-like Sensitivity (Sn), and their combined Sn-PPV accuracy (SPA) [22], also given in **S1 File** equation 9. In Sensitivity, the numerator is a sum of the maximal number of recalled nodes for each community and the denominator is a sum of the number of nodes in each community. Measures like these do not give equal importance to each of the known communities and assign higher values for recalling larger communities when compared to recalling smaller communities. For instance, an algorithm that perfectly recalls numerous smaller communities and does not recall much of a few bigger communities can get a worse sensitivity score when compared to an algorithm that does the opposite, *i.e.* recalls most of the big community and does not recall much of any of the smaller communities. Rather than inducing bias into a measure that decides which communities should be weighted higher, it may be a better idea to have a measure that gives equal weights to all communities. We define an unbiased sensitivity *Sn*_*u*_in **Fig 3e**, by dividing by the total number of known communities.

In PPV, the denominator sums, for each learned community, the sum of the subset of nodes in the learned community shared by all known communities. This does not contribute accurately to a precision-like measure, as nodes that are absent in known communities are ignored. For instance, a learned community that has all the nodes in a known community, but also includes a lot of possibly spurious nodes will be scored in the same way as a learned community which is an exact match to the known community. Further, in PPV, nodes in a learned community shared by multiple known communities get counted an extra number of times in the denominator. So if we share a set of nodes with multiple known communities we get penalized more than (i) if we share the set with only a few known communities, or (ii) if nodes of our community are shared with different known communities in a disjoint manner. The reasoning for allowing such behavior is again biased and does not support the detection of overlapping known communities. For example, a learned community that has a high overlap with 2 known communities (ex: a learned community with 10 nodes that shares all of its nodes with each of the known communities) will contribute lesser (0.5) to a PPV score than a learned community which overlaps lesser with one known community (ex: 6 nodes in a learned community with 10 nodes overlapping with only one known community, giving a 0.6 contribution to the PPV). To overcome these issues, we propose an unbiased PPV, *PPV*_*u*_ in **Fig 3e**, where we divide by the total number of learned communities. The corresponding unbiased accuracy is obtained by taking the geometric mean of the *PPV*_*u*_ and *Sn*_*u*_ as shown in **Fig 3e**.

From the sensitivity of measures plot in **Fig 3f**, we find that the FMMF score, the Qi et al F1 score (**S1 File** equation 8), and CMFF score are most sensitive to the pairwise overlap between communities, giving high values at the overlap coefficient yielding the best results, determined via visual inspection of the learned results, as follows. We observed highly overlapping, repetitive, and large numbers of similar learned protein complexes in our experiment on hu.MAP, such as several resembling the ribosome complexes at the high overlap threshold of 0.5 Jaccard coefficient, whereas, at low overlaps, we obtain a total small number of learned complexes, 84 learned complexes after removing proteins absent from known complexes. As we would like a high number of good quality complexes, we find that intermediate values of overlap Jaccard coefficient yield satisfactory results, for instance, at 0.25 Jaccard coefficient, we obtain 121 complexes after removing proteins absent from known complexes, with a high recall of known complexes and good observed quality, *i.e.* low numbers of very similar overlapping learned complexes. The clique-based measures from [4] - F-grand K-clique and F-weighted K-clique do not vary much with overlap, and the UnSPA, like the SPA, increases with increasing overlap threshold. However, the rate of increase of SPA w.r.t increasing overlap values is greater than UnSPA, yielding comparatively higher scores at undesirable high overlaps. In other words, instead of the desired decreasing trend from 0.25 to 0.5 Jaccard coefficient overlap, we have a highly increasing trend for SPA, compared to the almost constant trend for UnSPA - an improvement over SPA that can possibly be attributed to the unbiasing modification we have introduced. Therefore, for accurate evaluation in which redundancy (high overlap) is penalized, we recommend UnSPA over SPA, and primarily recommend the FMMF score, CMFF score, and the existing Qi et al F1 score (**S1 File** equation 8).

### Super.Complex applied to a human protein interaction network to detect protein complexes

#### Experiment details

We first test and ensure that the pipeline achieves perfect results on a toy dataset we construct comprising disconnected cliques of varying sizes, each corresponding to a known community, where we use all nodes as seeds for growth during the prediction step.

To learn potentially new human protein complexes, we apply Super.Complex on the human PPI network hu.MAP [4] using a community fitness function that is learned from known complexes in CORUM [3]. The network available on the website (http://hu1.proteincomplexes.org/static/downloads/pairsWprob.txt) has 7778 nodes and 56,712 edges, after an edge weight cutoff of 0.0025 was applied to the original 64,048 edges. There are 188 complexes after data cleaning, a set we term as ‘refined CORUM’, out of the original 2916 human CORUM complexes, which underscores the importance of minimizing any losses in the merging and splitting steps of the pipeline. In the data cleaning process, overlapping complexes with a Jaccard coefficient *j* greater than 0.6 are removed, as this value was used in the experiments of hu.MAP 2-stage clustering. Note that of the complexes from CORUM that were removed, there were over 1000 complexes that had fewer than 3 members, and the remaining removed complexes consisted of duplicates and disconnected complexes with edges from hu.MAP. Note, however, that hu.MAP was the highest confidence human protein interaction network integrating 3 large previous human protein-interaction networks, all built using high confidence data from large-scale (~9000) laboratory experiments. The edge weights of hu.MAP were trained using an SVM based on features obtained from experiments.

#### Experiment results

The best results, following different parameter sweeps from the experiment on hu.MAP are given in **Fig 4** with the best parameter values given in **Table 1**. From **Fig 4e**, we verify that the size distributions of the train and test sets are similar. In **Fig 4a**, we can see that we get a good precision-recall curve on the test set for the subgraph classification task as a positive or negative community, achieving an average precision score of 0.88 with a logistic regression model (which is the final ML model stacked on a set of other ML models and processors, output as the best model trained on the training set with 5-fold cross-validation and achieving a cross-validation score of 0.978).

**Fig 4.**
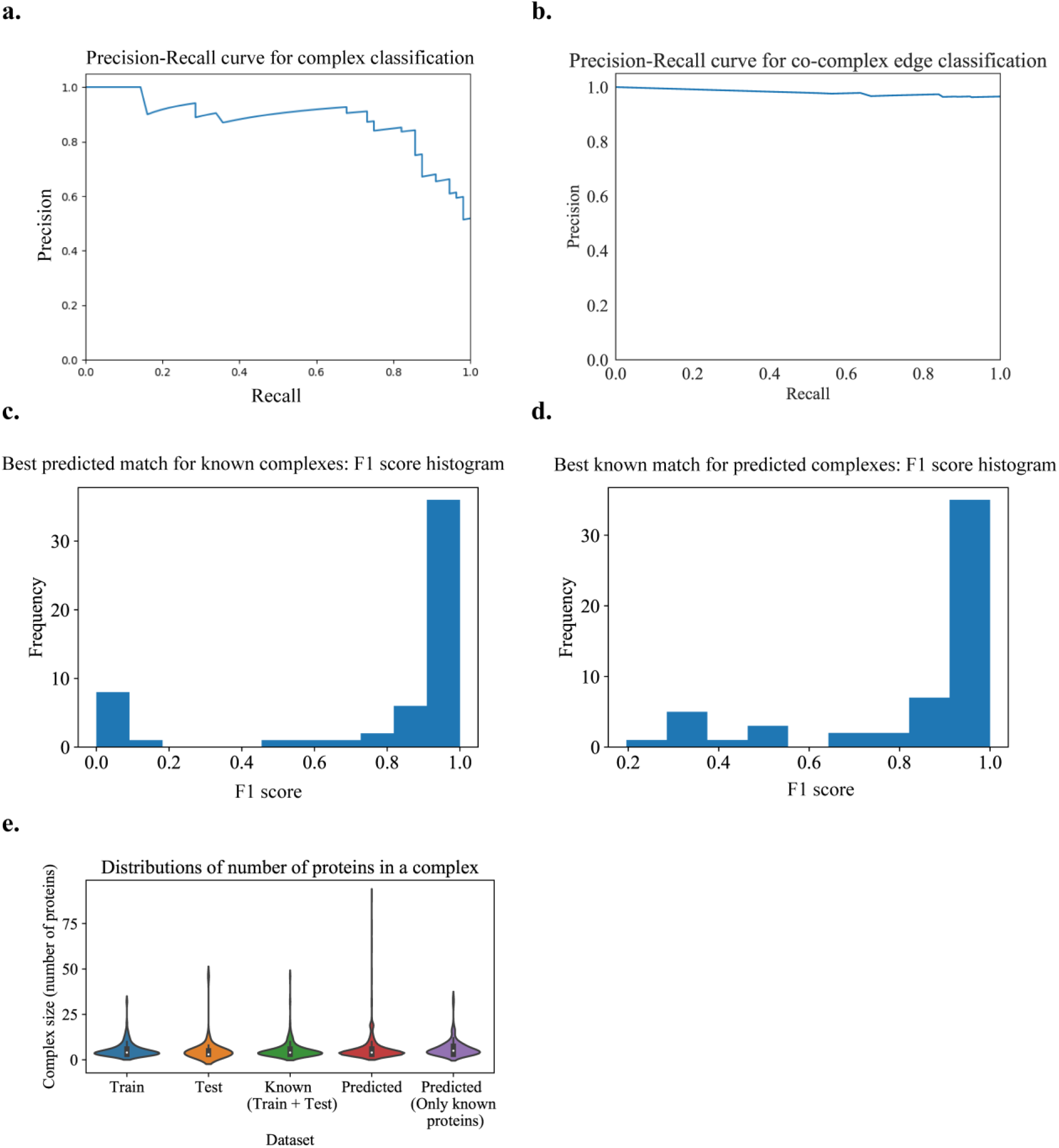
Learned human protein complexes with Super.Complex achieve good PR curves and follow similar size distributions as known complexes. **a.** PR curve for the best model (community fitness function) from the AutoML pipeline on the test dataset, for the task of classifying a subgraph as a community or not. **b.** Co-complex edge classification PR curve for final learned complexes. **c & d.** Best F-similarity score distributions per known complex and per learned complex. **e.** The size distributions of train, test, and all known complexes, learned complexes, and learned complexes after removing known complex proteins.

**Fig 5.**
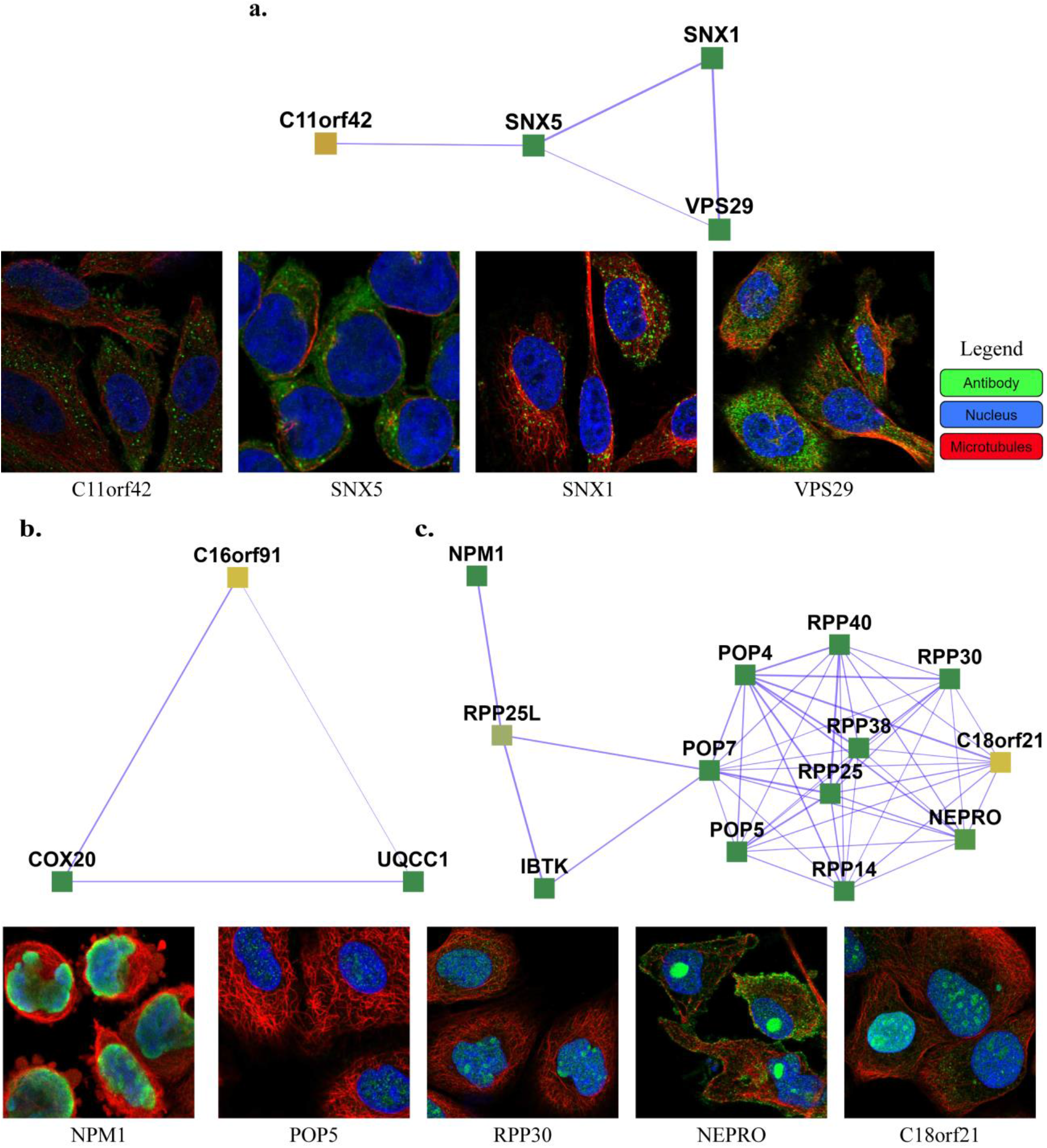
Examples of complexes with proteins having low annotation scores. **a.** C11orf42 constitutes the [Retromer complex](https://meghanapalukuri.github.io/Complexes/Predicted/Complex692.html) (SNX1, SNX2, VPS35, VPS29, VPS26A), potentially related to trafficking, with C11orf42 localized in cells to vesicles, similar to the other proteins of the complex (SNX1, SNX5, and VPS29) **b.** C16orf91 constitutes the [COX 20-C16orf91-UQCC1 complex](https://meghanapalukuri.github.io/Complexes/Predicted/Complex334.html), potentially localized to mitochondria like COX20. **c.** C18orf21 constitutes the [Rnase/Mrp complex](https://meghanapalukuri.github.io/Complexes/Predicted/Complex1012.html), with C18orf21, localized to nucleoli, closely interacting with nucleoplasm proteins of the complex such as RPP25, POP5, RPP14, [NEPRO](https://www.proteinatlas.org/ENSG00000163608-NEPRO/cell), RPP30, [IBTK](https://www.proteinatlas.org/ENSG00000005700-IBTK/cell), RPP25L, and NPM1. The images of subcellular localization are available from v20.1 of proteinatlas.org, as https://v20.proteinatlas.org/ENSG00000*/cell, where * is 180878-C11orf42, 028528-SNX1, 089006-SNX5, 111237-VPS29, 167272-POP5, 163608-NEPRO, 148688-RPP30, and 181163-NPM1. Note that localizations were measured in varying cell types, including HeLa, HEL, U2OS, and U-251 MG cells, across the highlighted proteins.

**Table 1.** Best parameters found and used in each of the experiments.

|  |  |  |  |  |
| --- | --- | --- | --- | --- |
| **PPI Network** | Hu.MAP | Yeast | Yeast | Yeast |
| **Experiment** | train: CORUM,<br>test: CORUM<br>(independent) | 1. train: TAP, test:<br>MIPS | 2. train: MIPS,<br>test: TAP | 3. train: MIPS, test:<br>MIPS |
| **Seeds** | All nodes | All nodes | All nodes | All nodes |
| **No. of negatives**<br>**sampled** | 10x positives | 1.1x positives | 1.1x positives | 1.1x positives |
| **Size (no. of nodes)**<br>**distribution for**<br>**negatives** | Uniform | Uniform | Uniform | Uniform |
| **Candidate sampling**<br>**Method** | ε- greedy +<br>iterative<br>simulated<br>annealing | ε- greedy +<br>iterative simulated<br>annealing | ε- greedy +<br>pseudo-metropolis | ε- greedy +<br>iterative simulated<br>annealing |
| **$\epsilon$** | 0.01 | 0.01 | 0.01 | 0.01 |
| **Sampling method**<br>**parameters** | T0 = 1.75 and $\alpha$<br>= 0.005 | T0 = 0.88 and $\alpha$ =<br>1.8 | Probability p =<br>0.1 | T0 = 0.88 and $\alpha$ =<br>1.8 |
| **No. of steps (specified or**<br>**inferred from known**<br>**complexes)** | 20 | 4 | 9 | 10 |
| **Neighbors considered**<br>**for growth** | All neighbors | All neighbors | All neighbors | All neighbors |
| **Merging method and**<br>**parameter** | Qi overlap<br>measure = 0.375 | Qi overlap measure<br>= 0.1 | Qi overlap<br>measure = 0.3 | Qi overlap measure<br>= 0.9 |

We use **Fig 4e** to set the maximum number of steps taken in the candidate complex growth stage as 20 and learn a total of 1028 complexes. On removal of non-gold standard proteins from these complexes for evaluation purposes, we obtain 131 complexes We get a good PR curve for the prediction of co-complex edges in comparison with known complex edges, as shown in **Fig 4b**. From **Fig 4c**, we can see that the best learned complex matches for known complexes have high F-similarity scores. Also, from **Fig 4d**, we can see that the best known complex matches for learned complexes have high F-similarity scores. Note that there may be unknown but true complexes that are learned by the algorithm that contribute to false positives.

In **Fig 4e**, we can see that learned complexes have a similar size distribution as known complexes. The small peak at size 20 is an artifact of our threshold on the maximum number of steps that can be taken in growing the complex. This means that either of our stopping criteria was not reached for these complexes, *i.e.* the criteria of a score less than 0.5 or no observed score improvement over a specified number of steps (here, 10).

Evaluation measures comparing learned complexes on hu.MAP by Super.Complex w.r.t known complexes from CORUM are given in **Tables 2 and S1 File Table 1**, along with the measures computed on the protein complexes comprising hu.MAP obtained from a 2 stage clustering method with the unsupervised ClusterONE algorithm applied first, followed by the unsupervised MCL algorithm. We observe that Super.Complex does better in terms of precision, as can be seen with the higher FMM precision value, while ClusterONE+MCL does better in terms of recall. This can be attributed to more number of complexes learned by ClusterONE+MCL (~4000 compared to ~1000 by Super.Complex) including a few highly overlapping complexes (the maximum pairwise overlap observed was 0.97 Jaccard coefficient), compared to the strict low overlap among complexes learned by Super.Complex (the maximum pairwise overlap observed was 0.36 Jaccard coefficient). We observe 4152 pairs of complexes learned by ClusterONE + MCL having an overlap greater than 0.36 Jaccard coefficient, the maximum pairwise overlap observed in learned complexes from Super.Complex. Note that while the values of F1 evaluation measures are similar, the results from ClusterONE+MCL were achieved by the authors after significant cross-validation, while Super.Complex was faster as detailed in **S1 File** Results (Performance).

**Table 2.** Evaluating learned complexes on hu.MAP w.r.t ‘refined CORUM’ complexes. Refined CORUM comprises 188 complexes after cleaning original CORUM complexes.

| Method | FMM |  |  | CMF<br>F1 score | Unbiased<br>Sn-PPV<br>accuracy | Qi et al<br>F1 Score<br>(t=0.5) | F-Grand<br>k-Clique | F-weighted<br>k-Clique |
| --- | --- | --- | --- | --- | --- | --- | --- | --- |
|  | Precision | Recall | F1 score |  |  |  |  |  |
| Super.Complex | **0.767** | 0.534 | **0.63** | 0.783 | 0.888 | 0.739 | **0.785** | **0.972** |
| hu.MAP<br>(ClusterONE + MCL) | 0.471 | **0.686** | 0.559 | **0.797** | **0.911** | **0.764** | 0.77 | 0.967 |

### State of the art comparison: Super.Complex achieves good evaluation measures and performance

To compare our method with published results from existing methods, we perform experiments on the data used by these methods in their experiments - a yeast PPI network, DIP - Database of Interacting Proteins [23] with known protein complexes from MIPS - Munich Information Center for Protein Sequence [24] and TAP--Tandem Affinity Purification [25]. Specifically, for an accurate comparison, we use the same PPI network (projection of DIP yeast PPI network on MIPS + TAP proteins) and known protein complexes, available from the ClusterEPs software website. The results from **Table 3** show that our method outperforms all 6 supervised as well as 4 unsupervised methods (by achieving the highest F1 score and precision values) in the yeast experiments. Specifically, Super.Complex achieves the highest F1 score value (87% higher on average, 63% higher by median) when compared to the 10 other methods, highest precision value (110% higher on average, 72% higher by median) when compared to the 10 other methods, higher recall (92 % higher on average, 45% higher by median) when compared to 8 other algorithms with lower recall values (30% lower on average and by median) when compared to only 2 methods (ClusterSS and ClusterEPs, considered next best as per the F1 score, a metric which gives a better notion of the performance of an algorithm than just the recall or precision measure taken alone). When comparing with the 2 algorithms where Super.Complex has lower recall, it makes up for this by significantly outperforming the precision measure (55% higher on average and by median) to achieve higher F-1 scores (12% higher on average and 14% higher by median). Also, as we have noted earlier, for this application of detecting protein complexes, validation of results usually involves time-taking and expensive biological experiments, therefore, an algorithm like Super.Complex yielding a low number of false positives (translating to high precision) is more desirable (even with lower recall) than an algorithm that is able to identify many existing communities but with high false positive rates (translating to higher recall but low precision). From **Table 3**, similar to observations of metrics from the experiments on hu.MAP in **Tables 2** and **S1 File Table 1**, we obtain high precision values with Super.Complex, suggesting that many of the learned protein complexes are of high quality. On performance, we discuss the time complexity of Super.Complex in the **S1 File** Methods (Time complexity). The whole pipeline was completed in an order of minutes with Super.Complex (including the AutoML step executed on a single Skylake compute node, along with parameter-sweeps for the candidate community sampling step executed on 4 Skylake compute nodes - each with 48 cores @2GHz clock rate). We attempted to run other algorithms on hu.MAP as well, but were unsuccessful due to unavailability of code or limited scalability, as detailed in **S1 File** Results (SOTA availability).

**Table 3.** Comparing our method with 6 supervised and 4 unsupervised methods on a yeast PPI network. Precision, recall, and F-measures are from Qi et al. Parameters for each of the Super.Complex experiments are given in **Table 1**.

|  | **Train** | **Test** | **Precision** | **Recall** | **F-measure** |
| --- | --- | --- | --- | --- | --- |
| **Super.Complex** | TAP | MIPS | **0.841** | 0.629 | **0.72** |
| ClusterSS | TAP | MIPS | 0.526 | 0.807 | 0.636 |
| ClusterEPs | TAP | MIPS | 0.606 | 0.664 | 0.633 |
| RM | TAP | MIPS | 0.489 | 0.525 | 0.506 |
| SCI-BN | TAP | MIPS | 0.219 | 0.537 | 0.312 |
| SCI-SVM | TAP | MIPS | 0.176 | 0.379 | 0.240 |
| ClusterONE |  | MIPS | 0.428 | 0.435 | 0.431 |
| COACH |  | MIPS | 0.364 | 0.495 | 0.419 |
| CMC |  | MIPS | 0.46 | 0.38 | 0.416 |
| MCODE |  | MIPS | 0.4 | 0.1 | 0.16 |
| **Super.Complex** | MIPS | TAP | **0.718** | 0.581 | **0.642** |
| ClusterSS | MIPS | TAP | 0.477 | 0.864 | 0.614 |
| ClusterEPs | MIPS | TAP | 0.424 | 0.782 | 0.548 |
| RM | MIPS | TAP | 0.424 | 0.433 | 0.429 |
| SCI-BN | MIPS | TAP | 0.312 | 0.489 | 0.381 |
| SCI-SVM | MIPS | TAP | 0.247 | 0.377 | 0.298 |
| ClusterONE |  | TAP | 0.480 | 0.46 | 0.47 |
| COACH |  | TAP | 0.387 | 0.533 | 0.449 |
| CMC |  | TAP | 0.447 | 0.353 | 0.395 |
| MCODE |  | TAP | 0.422 | 0.127 | 0.195 |
| **Super.Complex** | MIPS | MIPS | **0.552** | **0.733** | **0.63** |
| NN | MIPS | MIPS | 0.333 | 0.491 | 0.397 |

### Learned human protein complexes from Super.Complex, and applications to COVID-19 and characterizing unknown proteins

We provide interactive lists and visualizations of the <u>1028 learned human protein</u> <u>complexes</u> by Super.Complex, along with <u>refined</u> and <u>original</u> CORUM complexes as a resource on https://sites.google.com/view/supercomplex/super-complex-v3-0. The high precision values obtained by Super.Complex in **Table 2** suggest that many of the learned complexes are of high quality, since the ones with proteins from known complexes match individual known complexes closely. We provide individual community fitness function scores for each of the learned complexes, and rank the list of learned complexes by this score to help identify good candidates for investigation for various applications. In this section, we analyze learned human protein complexes by Super.Complex, aiming to provide easily accessible resources for two biological applications that can be investigated further by researchers in the future. We highlight learned complexes with uncharacterized proteins to provide experimental candidates for functional characterization. In the second application, we construct an interactive map of SARS-CoV-2 protein interactions with 234 learned human protein complexes from Super.Complex using protein-interaction information between SARS-CoV-2 proteins and human proteins [26]. We also provide a list of complexes interacting with SARS-CoV-2 proteins ranked by their possible importance, which can be used to determine potential COVID-19 drug targets (S1 Fig & **S1 File** Results (SARS-CoV-2 affected protein complexes)).

#### Uncharacterized proteins and their complexes

111 uncharacterized proteins (Uniprot [27] annotation score unknown or less than 3) and their corresponding learned 103 complexes are presented on the website (https://meghanapalukuri.github.io/Complexes/* where * is Protein2complex_annotated.html and Complex2proteins_annotated.html). Three examples of uncharacterized proteins (C11orf42, C18orf21, and C16orf91) along with their corresponding complexes are highlighted in **Fig 6**. C11orf42 could potentially be related to trafficking, as it is a part of a complex with 30% similarity to the retromer complex, (*i.e.* with 0.3 Jaccard similarity to the known CORUM retromer complex), with additional evidence available from the Human Protein Atlas (HPA) [28] (available from http://www.proteinatlas.org) showing subcellular localization to vesicles, similar to other proteins of the complex. C18orf21 also has evidence from HPA, localized to the nucleoli and interacting with other proteins of a complex with 50% similarity to the Rnase/Mrp complex with most members in the nucleoli/nucleoplasm. Further evidence from [29] also independently supports C18orf21 as a cellular component of the ribonuclease MRP complex and a participant in ribonuclease P RNA binding as it exhibits significant co-essentiality across cancer cell lines with the POP4, POP5, POP7, RPP30, RPP38, and RPP40 proteins. C16orf91 could potentially be localized to mitochondria like other proteins of the COX 20-C16orf91-UQCC1 complex, with independent experimental evidence from [30].

## Data and Code Availability

We make interactive visualizations of our learned protein complexes freely available as a resource at https://sites.google.com/view/supercomplex/super-complex-v3-0, which includes downloadable sets of interactions and complexes, including the 234 complexes that are potentially linked to COVID-19 and SARS-CoV-2 infection, and the set of 111 uncharacterized proteins implicated in 103 complexes. Our code is available at https://github.com/marcottelab/super.complex. To simplify reanalysis, the full interactome datasets are additionally deposited in Zenodo, DOI: http://doi.org/10.5281/zenodo.4814944

## Acknowledgments

The authors gratefully acknowledge Benjamin Liebeskind for a TPOT code wrapper, Kevin Drew for computing some previous evaluation measures, Claire McWhite for critical reading of the manuscript, and funding from the Welch Foundation (F-1515) and National Institutes of Health (R35 GM122480, R01 HD085901, R21 HD103588) to E.M.M. The funders’ websites are https://www.welch1.org/ and https://www.nih.gov/ respectively. The funders had no role in study design, data collection and analysis, decision to publish, or preparation of the manuscript.

## Supplementary Information

**S1 File. Document containing supplementary tables, results and methods.**

a. **Table 1**. Comparing Super.Complex with 2-stage clustering on the hu.MAP dataset using 6 existing and 3 new evaluation metrics shows comparable performance for both algorithms.
b. **Results**: Algorithm Guarantees, Robustness of the Super.Complex algorithm, Performance, SOTA Availability, and SARS-CoV-2 affected protein complexes.
c. **Methods**: Topological Features, Similarity measures for evaluation, Evaluation with existing measures, Time complexity

**S1 Fig. SARS-CoV-2 - human protein complex map showing complexes identified by Super.Complex. a.** A section of the full map, featuring SARS-CoV-2 nsp4 and orf6 and their interacting human protein complexes **b.** A protein complex with a 30% match to the Nup 107-160 subcomplex interacts with both SARS-CoV-2 nsp4 and orf6 **c.** Map of SARS-CoV-2 nsp2 interactions with human proteins and their corresponding complexes **d.** A complex with a 20% match to the Endosomal targeting complex, and **e.** A complex with a 40% match to the retromer complex, both of which interact with SARS-CoV-2 nsp2. An interactive map is available at https://meghanapalukuri.github.io/Complexes/SARS_COV2_Map_only_mapped_complexes_names.html.

## Notes

https://doi.org/10.5281/zenodo.4814944

https://github.com/marcottelab/super.complex

https://sites.google.com/view/supercomplex/super-complex-v3-0

### Competing Interest Statement

The authors have declared no competing interest.

### Summary of Updates

Supplementary file added with new analyses and discussion also referenced in the main paper; Figure 2 revised; Paper re-organized with methods section before results; Some sections in main paper moved to supplementary; Some changes in the main paper are: 1. Addition of Table 1 replacing text descriptions of parameters used in experiments. 2. More consistent naming of steps in the Super.Complex pipeline and 3. quantifying better the difference in performance of Super.Complex and SOTA algorithms. In the supplementary, we add a table comparing Super.Complex performance with more evaluation metrics, a section on robustness of the Super.Complex algorithm and time complexity.

