## Supplementary_S1_File for "Super.Complex: A supervised machine learning pipeline for molecular complex detection in protein-interaction networks"

### Supplementary Tables

**Table 1. Comparing Super.Complex with 2-stage clustering on the hu.MAP dataset using 6 existing and 3 new evaluation metrics shows comparable performance for both algorithms.** In summary, Super.Complex achieves better performance on 7 precision-like metrics (18% higher on average, 9% by median with 2 metrics doing better with 2-stage clustering by 3% on average and by median), while the 2-stage clustering method achieves better performance on recall measure on all 9 metrics by 10% on average and 9% by median. Super.Complex achieves better F1-score-like measures on 4 of the metrics by 4% on average and 2% by median, compared to 5 wins by 2-stage clustering by 2% on average and by median.

| **Metric** | | | Super.Complex | hu.MAP (ClusterONE + MCL) |
|---|---|---|---|---|
| No. of learned protein complexes | | | 1028 | 4659 |
| No. of learned protein complexes after removing non-gold standard proteins | | | 131 | 274 |
| New metrics | FMM | Precision | **0.767** | 0.471 |
| | | Recall | 0.534 | **0.686** |
| | | F1 score | **0.63** | 0.559 |
| | CMF | Precision | **0.839** | 0.804 |
| | | Recall | 0.733 | **0.789** |
| | | F1 score | 0.783 | **0.797** |
| | UnSPA | $PPV_u$ | 0.903 | **0.93** |
| | | $Sn_u$ | 0.874 | **0.893** |
| | | UnSPA | 0.888 | **0.911** |
| Existing metrics | SPA | PPV | 0.885 | **0.907** |
| | | Sn | 0.675 | **0.681** |
| | | SPA | 0.773 | **0.786** |
| | Qi et al | Precision | **0.832** | 0.763 |
| | | Recall | 0.665 | **0.766** |
| | | F1 score | 0.739 | **0.764** |
| | K-clique | Precision | **0.992** | 0.825 |
| | | Recall | 0.674 | **0.735** |
| | | F1 score | **0.785** | 0.772 |
| | Weighted K-clique | Precision | **0.998** | 0.973 |
| | | Recall | 0.948 | **0.962** |
| | | F1 score | **0.972** | 0.968 |
| | MMR | Precision | **0.714** | 0.616 |
| | | Recall | 0.664 | **0.747** |
| | | F1 score | **0.688** | 0.675 |

| | | | | |
|---|---|---|---|---|
| | PR | Precision | **0.738** | 0.681 |
| | | Recall | 0.692 | **0.759** |
| | | PR product | 0.714 | **0.718** |

### Supplementary Results

#### Algorithm Guarantees

The train and test communities are guaranteed to be independent by the proposed splitting algorithm. Due to the seeding and growth heuristics we employed, the learned communities are guaranteed to be internally connected at every step of the iteration. With an overlap threshold t, the merging algorithm guarantees that no 2 learned communities have an overlap greater than or equal to t. If an overlap threshold of 0 is provided, the merging algorithm yields disjoint, non-overlapping learned communities which guarantee separation, *i.e.* no pair of learned communities can be merged to yield a community of higher community score. The number of maximum iterations provided guarantees that the seeding and growth converge to a solution.

#### Robustness of the Super.Complex algorithm

In the candidate community search, we can expect that the learned complexes will differ from experiment to experiment due to 3 modes of possible stochasticity namely, (i) the $\epsilon$-greedy heuristic, (ii) the pseudo-metropolis or ISA heuristic, and (iii) the post-processing algorithm merging pairs of highly overlapping communities. A re-run of the search with the same parameters as the best results on hu.MAP yielded 1025 complexes, compared to 1028 in the original experiment run. Of these, 999 were identical. We compare the sets of learned complexes from both runs and observe high correlation, as observed from an FMM F1 score of 0.984, CMF F1 score of 0.988, unbiased accuracy of 0.989 and a Qi et al F1 score of 0.985. This indicates that despite the stochasticity, we can expect fairly stable and robust results.

#### Performance

The current problem of learning the best community fitness function and growing multiple seeds on a network into candidate communities poses an interesting design challenge. Multiple architectures are valid and can be the best based on the constraints of the computing systems. Our design performs well and is capable of giving results in an order of minutes for a network with a similar scale (~10k nodes, ~100k edges) as hu.MAP. With default parameters, a single run of the pipeline on hu.MAP gave reasonably good results in around 30 minutes, on 4 Skylake nodes (each with 48 cores @2GHz clock rate). In more detail, it averages ~10 min per parameter set of the candidate community sampling part of the pipeline using 4 Skylake nodes. For the best results, we ran multiple parameter sweeps for a few hours (a total of around 100 sets), achieving only a minor improvement in results. The AutoML model can yield good models in an order of minutes as well, however, we ran it for a few hours on a single node to obtain the best model. For accessibility to run on non-HPC (high-performance computing) systems as well, using the optimizations provided and setting low thresholds, it is possible to run the pipeline serially in a matter of hours, even on a personal laptop. A discussion on time complexity is included in Supplementary Methods (Time Complexity).

#### SOTA Availability

We performed experiments with the implementation of the state-of-the-art method SCI-SVM on hu.MAP whose Matlab code was available and which we were able to run on a single core of a Skylake node (2GHz clock rate). The implementation with the default parameters on a toy dataset and a human dataset resulted in the identification of a large number of small complexes and a very few big complexes, despite sufficient training with big complexes. As big complexes were only a few, the results have low precision and recall. Computed precision, recall, and F1 scores as per equation 10 for even a low threshold $t$ of 0.2 are 0.016, 0.0075, and 0.0103 respectively for hu.MAP. A single run of SCI-SVM took around 3 hours to complete and yield these results, unlike better results yielded by Super.Complex in around 30min. Additional runs tuning different parameters in SCI-SVM did not improve results.

In our experiments, we were unable to get the UI (User Interface) for the ClusterEPs software to complete execution on a personal laptop for hu.MAP. Note that Super.Complex on the other hand finishes running on the same personal laptop and provides results after a few hours. Code was not available for ClusterEPs, or the other supervised methods in **Table 3** – RM, NN and ClusterSS.

#### SARS-CoV-2 affected protein complexes

SARS-CoV-2 is the novel coronavirus that resulted in the pandemic which started at the end of 2019. The virus infects human cells through a spike protein and enters the cell, and subsequently, the proteins of the virus have the potential to interact with multiple human proteins. We preliminarily investigated the human protein complexes that may be affected by the virus, and thus contribute, directly or indirectly to COVID-19, the disease caused by SARS-CoV-2. This study was made possible by timely work identifying 332 human proteins that physically interact with SARS-CoV-2 proteins, discovered using AP-MS experiments [3].

We consider a protein complex to be linked to COVID-19 if any of the proteins in the human complex interact with a SARS-CoV-2 protein, *i.e.*, if the human protein complex contains one or more of the above mentioned 332 human proteins. Super.Complex learns 234 complexes linked to COVID-19, all except one new, *i.e.* not perfectly matching a known CORUM complex. While CORUM contains 430 complexes linked to COVID-19 ('original CORUM'), we obtain 214 complexes after cleaning ('refined CORUM'), *i.e.* removing complexes with sizes lesser than or equal to 2 and using the merging algorithm described in the Materials and Methods section to ensure no more than 50% Jaccard overlap between any two SARS-CoV-2 complexes. All SARS-COV2 linked protein complexes from Super.Complex, scored based on the number of proteins in a complex interacting with a SARS-CoV-2 protein, along with lists of refined CORUM and original CORUM complexes linked to SARS-COV2 are available on the website (https://meghanapalukuri.github.io/Complexes/* where * is Complex2proteins_covid.html, CORUM_Complex2proteins.html, and originalCORUM_Complex2proteins_covid.html respectively). **S1 Fig** shows examples of human protein complexes likely especially relevant to the SARS-CoV-2 life cycle, as they involve multiple interactions from SARS-CoV-2 proteins and members of the same (or a functionally related) complex. Visualizing the SARS-CoV-2 interacting proteins in terms of their native assemblies serves to emphasize the preferential interaction of SARS-CoV-2 proteins with certain cellular systems, as for nsp4 and orf6 both interacting with the nuclear pore complex, and may help highlight important aspects of the SARS-CoV-2 life cycle for future study.

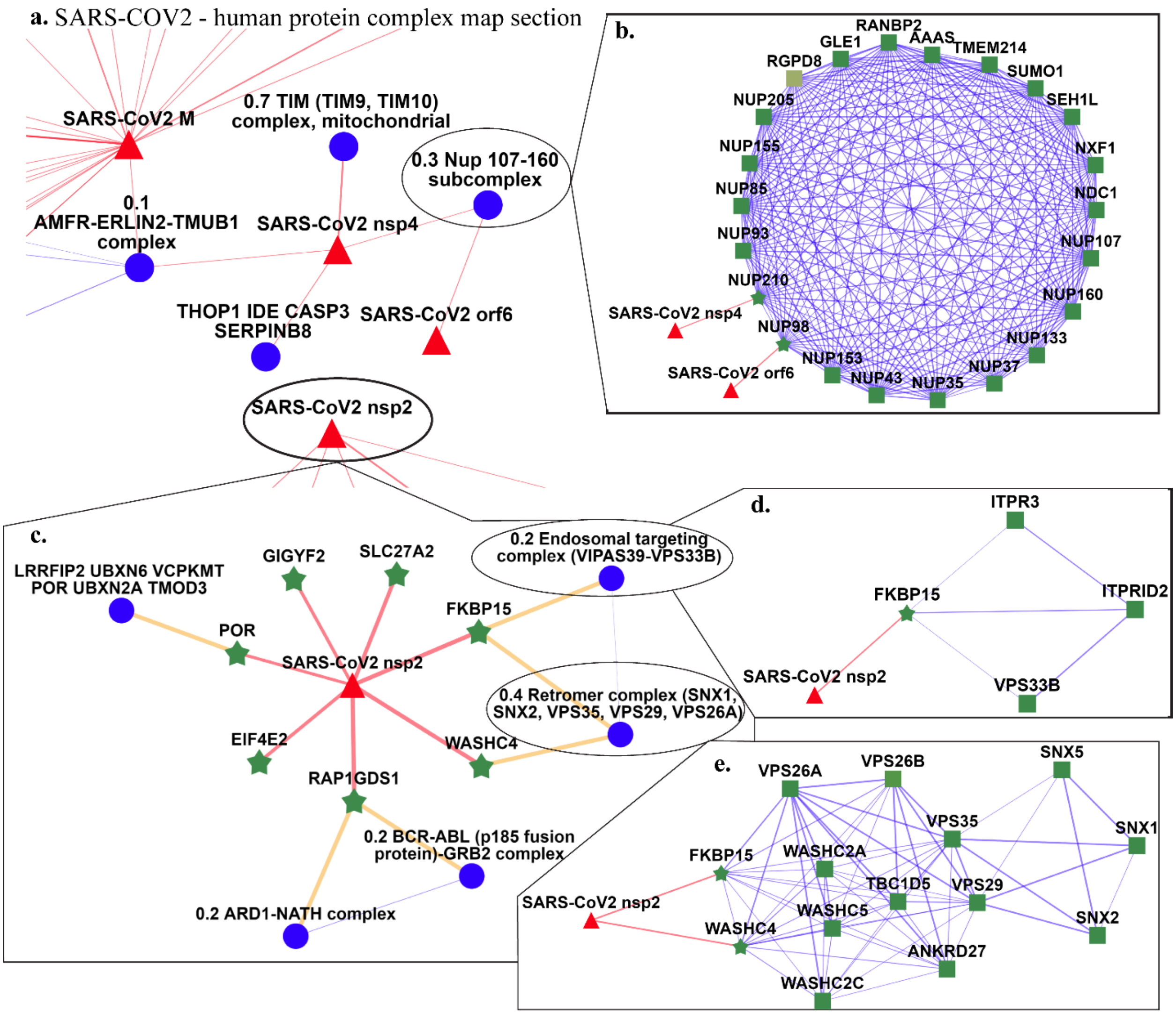


**S1 Fig. SARS-CoV-2 - human protein complex map showing complexes identified by Super.Complex. a.** A section of the full map, featuring SARS-CoV-2 nsp4 and orf6 and their interacting human protein complexes **b.** A protein complex with a 30% match to the Nup 107-160 subcomplex interacts with both SARS-CoV-2 nsp4 and orf6 **c.** Map of SARS-CoV-2 nsp2 interactions with human proteins and their corresponding complexes **d.** A complex with a 20% match to the Endosomal targeting complex, and **e.** A complex with a 40% match to the retromer complex, both of which interact with SARS-CoV-2 nsp2. An interactive map is available at https://meghanapalukuri.github.io/Complexes/SARS_COV2_Map_only_mapped_complexes_names.html
.

### Supplementary Methods

#### Topological Features

These features include 5 subgraph specific features - the no. of nodes in the subgraph, the 1st 3 singular values of the adjacency matrix of the subgraph, and the weighted intra-cluster density, obtained by using the weighted sum of edges instead of the number of edges in the numerator of the intra-cluster density defined in equation 2. The remaining 13 features are statistical measures of the combinations of features defined for individual nodes of the subgraph. We use the median, mean, variance, and maximum value of the degree $d_v$of a vertex *v*, which is the sum of the weights of the edges connecting the node to its immediate neighbors. Next, we use the mean, variance, and maximum value of the clustering coefficient of a vertex, $c_v$, which gives the ratio of the number of triangles it is a part of considering only its first nearest neighbors and the number of all such possible triangles, *i.e.*,

$$c_v = \frac{No.of\ edges\ connecting\ pairs\ of\ the\ neighbors\ of\ v}{No.of\ all\ possible\ edges\ connecting\ pairs\ of\ neighbors\ of\ v} = \frac{|\{(v',v'') \mid v',v'' \in N(v), v' \in N(v'')\}|/2}{|N(v)|^* (|N(v)|-1)/2} \quad (1)$$

Where, *N(v)* is the set of neighbors of vertex v. We use the mean, variance, and maximum value of the degree correlation of a node *v*, which is the average degree of all neighbors of node *v*, *i.e.*,

$$DC_v = \frac{\sum_{v' \in N(v)} d_{v'}}{|N(v)|} \quad (2)$$

The last 3 features are the mean, variance, and maximum value of the weights of edges of the subgraph.

#### Similarity measures for evaluation

For evaluation, a similarity measure is used to compare two communities $C_1$ and $C_2$ and they are determined to be a match if the similarity measure value for the two communities is above a threshold. Some similarity measures include:

$$\text{Jaccard similarity}(C_1, C_2) = \frac{|C_1 \cap C_2|}{|C_1 \cup C_2|} \quad (3)$$

$$Neighborhood\ affinity = \frac{|C_1 \cap C_2|^2}{|C_1| * |C_2|} \quad (4)$$

Another similarity measure which requires a precision-like and recall-like component to both exceed a threshold is given by:

$$\text{Qi overlap measure: } \frac{|C_1 \cap C_2|}{|C_1|} > t \text{ and } \frac{|C_1 \cap C_2|}{|C_2|} > t \quad (5)$$

where *t* is the user-specified threshold, usually set to 0.5

Here, $|C|$ is the number of nodes in community *C*, $|C_1 \cap C_2|$ is the number of nodes common to both communities, and $|C_1 \cup C_2|$ is their union, *i.e.* the total number of nodes in the two communities with each node counted once.

Once we have a way to match a learned community to a known community or vice-versa using a similarity measure, standard machine learning metrics such as precision, recall, and F1 scores can be computed.

$$Qi\ et\ al\ Precision\ p = \frac{No.\ of\ learned\ communities\ that\ match\ known\ communities}{No.\ of\ learned\ communities} \quad (6)$$

$$Qi\ et\ al\ Recall\ r = \frac{No.\ of\ known\ communities\ that\ match\ learned\ communities}{No.\ of\ known\ communities} \quad (7)$$

$$Qi\ et\ al\ F1\ score = \frac{2pr}{(p+r)} \quad (8)$$

#### Evaluation with existing measures

To compare the set of learned communities with known communities, we first construct a set of reduced communities containing nodes only present in known communities. We retain communities with 3 or more nodes only. We use a plethora of evaluation measures, including the proposed measures detailed in the results section, along with existing measures such as S1 equation 5 based Qi et al precision, recall and F1 score (S1 equations 6-8), sensitivity. PPV, their associated accuracy, and MMR. The Qi et al precision, recall, and F1 score consider one-to-many associations, *i.e.* a learned community can be matched to multiple known communities and vice versa. One-to-one matches are made by measures such as the MMR - maximum matching ratio which calculates the weighted recall (the weights here are neighborhood affinity similarity scores) after selecting a set of one to one matches such that the sum of their similarity scores is maximal. Implicit one-to-one associations are made by measures such as sensitivity (Sn) and positive predictive value (PPV) which resemble a recall-like measure and a precision-like measure respectively. The implicit one-to-one associations made here correspond to picking the community that best matches another community in terms of the number of common nodes. The Sn-PPV accuracy (SPA) combines sensitivity and PPV, resembling a composite measure like the F1 score that combines Precision and Recall. Note that this measure makes one-to-many matches. Letting the set of learned communities be *L* and the set of known communities be *K*, we have,

$$Sn = \frac{\sum_{C_k \epsilon K} \max_{C_l \epsilon L} | C_l \cap C_k |}{\sum_{C_k \epsilon K} | Ck |},$$

$$PPV = \frac{\sum_{C_l \epsilon L} \max_{C_k \epsilon K} | C_l \cap C_k |}{\sum_{C_l \epsilon L} \sum_{C_k \epsilon K} | C_l \cap C_k |}, SPA = \sqrt{(Sn^* PPV)} \quad (9)$$

We also plot PR curves for edges learned. The learned communities are evaluated against all known communities, only training communities, and only testing communities. We report the results on the test communities for the yeast experiment in **Table 3**.

#### Time complexity

The average time complexity of the algorithm is determined by the two main steps - training the AutoML algorithm TPOT and the candidate community search step. The time complexity of TPOT is not explicitly mentioned in their papers [4-6], but from the method description, in our use, we can make a simple estimate. It would be at least $O\left(\frac{g \times p \times m \times s \times f}{c}\right)$, where $g$ is the number of generations, $p$ is the population size, $m$ is the number of machine learning models and feature preprocessor types tried, $s$ is the number of samples, $f$ is the number of features used, and $c$ is the number of processes in the single compute node running the AutoML step.

The candidate community search step has a complexity of $O\left(\frac{X \times G^4 \times K \times S}{P}\right)$, keeping in mind that in the worst case, the feature extraction step has a complexity of $O(n^3)$ for a subgraph with $n$ nodes. $P$ is the total number of processes used to run this code in parallel, $S$ is the number of seeds used to sample candidate communities which is typically $N$, the number of nodes in the network, $G$ is the number of steps for each community's growth and is also taken to be the final number of nodes in the subgraph after growth for complexity calculation (as both are similar), $K$ is the average degree of the graph. $X$ is the machine learning model inference time for one value, which can be assumed a small constant here in cases with fairly simple models. We note that in an alternate method that we provide, one can specify the number of neighbors to check, say $M$ instead of checking all neighbors, achieving gains when $M < K$, with the complexity now as $O\left(\frac{X \times G^4 \times M \times S}{P}\right)$. Using all nodes as seeds ($S = N$), in very large sparse protein interaction networks (small degree $K \ll N$) with small complexes (for example, we use $G < 10$ nodes in our yeast experiments) and considering that the machine learning model inference time is constant, the complexity can be considered $\sim O\left(\frac{N}{P}\right)$. In problems where communities are large making the polynomial greater than the number of nodes (*i.e.*, $G^4 > N$) time complexity will be determined instead by the community size $G$. For comparison, the authors note that SCI-SVM and SCI-BN have a complexity (in our notation) of $O(G^5 \times K \times S)$, while other supervised methods do not mention their time complexities. We note that all the other supervised algorithms in the best case scenario, assuming efficient implementation of the growth strategy for a single node would still have a complexity of $O(N)$ due to the serial nature of the algorithms, compared to our best case scenario of $O\left(\frac{N}{P}\right)$, where one can achieve gains by increasing the number of parallel processes $P$.
